# Regulation of the JMJD3 (KDM6B) histone demethylase in glioblastoma stem cells by STAT3

**DOI:** 10.1101/063040

**Authors:** Maureen M. Sherry-Lynes, Sejuti Sengupta, Shreya Kulkarni, Brent H. Cochran

## Abstract

The growth factor and cytokine regulated transcription factor STAT3 is required for the self-renewal of several stem cell types including tumor stem cells from glioblastoma. Here we show that STAT3 inhibition leads to the upregulation of the histone H3K27me2/3 demethylase Jmjd3 (KDM6B), which can reverse polycomb complex-mediated repression of tissue specific genes. STAT3 binds to the Jmjd3 promoter, suggesting that Jmjd3 is a direct target of STAT3. Overexpression of Jmjd3 slows glioblastoma stem cell growth and neurosphere formation, whereas knockdown of Jmjd3 rescues the STAT3 inhibitor-induced neurosphere formation defect. Consistent with this observation, STAT3 inhibition leads to histone H3K27 demethylation of neural differentiation genes, such as Myt1, FGF21, and GDF15. These results demonstrate that the regulation of Jmjd3 by STAT3 maintains repression of differentiation specific genes and is therefore important for the maintenance of self-renewal of normal neural and glioblastoma stem cells.

## INTRODUCTION

Glioblastoma stem cells (GBM-SC) are highly tumorigenic, and share many properties with normal neural stem cells [1, 2]. GBM-SC have gene expression patterns that more closely resemble their tumor of origin than do matched serum-derived cell lines [1]. STAT3 is a transcription factor that is activated by many cytokines and growth factors, and has a demonstrated role in oncogenesis of many human tumors including glioblastoma [3, 4]. STAT3 is required for the maintenance of pluripotency of murine embryonic and neural stem cells and facilitates reprogramming of somatic cells to the pluripotent state [5–7].

We and others have previously shown that the transcription factor STAT3 is essential for glioblastoma stem cell proliferation and multipotency [8–11]. Inhibition or RNAi knockdown of STAT3 leads to a dramatic decrease in proliferation and neurosphere formation, as well as loss of stem cell markers [11]. Interestingly, this phenotype is irreversible. Transient treatment with STAT3 inhibitors for as little as four hours leads to a permanent loss of neurosphere formation capacity, despite the fact that STAT3 signaling is restored upon drug removal [11]. This observation suggests that STAT3 regulates the epigenetic state of the cells, thereby causing a stable change in the ability of the cells to respond to stem cell growth factors.

In stem cells, including normal neural stem cells and GBM-SC, polycomb-mediated repression of differentiation specific genes is a major mechanism by which multipotency is maintained [12]. PRC2 adds methyl groups to histone H3K27, which leads to the recruitment of the PRC1 and the heritable inhibition of transcription [13, 14]. The H3K27me2/3 histone demethylase Jmjd3 (KDM6B) antagonizes the enzymatic activity of the polycomb repressive complex 2 (PRC2) [15–18]. Jmjd3 demethylates histone H3K27 at the promoters of neuronal-specific genes in mice [19], and is required for neural differentiation of murine embryonic stem cells and for proper neural differentiation in adult mice[20, 21]. Jmjd3 expression can also be rapidly induced in macrophages in response to LPS stimulation [16].

It is becoming increasingly apparent that H3K27 trimethylation is aberrantly regulated in several cancers. Inactivating mutations have been identified in the H3K27 demethylase *UTX(KDM6A)*, suggesting that UTX acts as a tumor suppressor [22]. Overexpression of EZH2 has been demonstrated in multiple cancer types as well, including medulloblastoma and glioblastoma brain tumors [23–25]. EZH2 expression is necessary for the self-renewal of glioblastoma stem cells [24]. Mutations in histone H3 itself have been found in pediatric glioblastoma [26–28]. Recently, it has been shown that Jmjd3 is induced during glioblastoma stem cell differentiation and is mutated in some GBM-SC and that constitutive Jmjd3 expression in GBM-SC inhibits their tumorigenesis [29]. These findings suggest that the aberrant maintenance of H3K27 methylation contributes to oncogenesis. Based on these observations and the STAT3 inhibition phenotype, we investigated the possibility that the H3K27 demethylase Jmjd3 may mediate effects of STAT3 in glioblastoma stem cells.

## MATERIALS AND METHODS

### Culture of glioblastoma stem cells and normal neural stem cells

Glioblastoma stem cell lines and culture conditions were described previously [11]. Normal human neural stem cells derived from H9 embryonic stem cells were obtained from Gibco (Carlsbad, CA). Cells were grown according to instructions of the manufacturer. For neurosphere formation assays, cells were dissociated from the coated plates with accutase and plated in neural stem cell media in uncoated plates. STAT3 inhibitors S3I-201, STA-21 (NCI/DTP Open Chemical Repository) or BP1-102 (EMD Millipore, Billerica, MA) were used at the indicated concentrations. Recombinant BMP4 (Life technologies, Grand Island, NY) was added to the culture media when indicated.

### Immunoblotting

Immunoblots were performed as previously described [11] with the exception that immunoblots to be probed with anti-Jmjd3 antibody were transferred in 10 mM CAPS (3-[Cyclohexylamino]-1-propanesulfonic acid, pH 10.5) buffer. Anti-β-Actin (1:2000) was obtained from Sigma (St. Louis, MO); anti-STAT3 (1:1000), anti pSer-STAT3 (1:200), anti-pTyr STAT3 (1:500) were obtained from Cell Signaling Technologies (Beverly, MA). Anti-Jmjd3 was obtained from Abgent (San Diego, CA) or generously provided by the lab of Yang Shi.

### RT-qPCR

RT-qPCR was performed as previously described [11]. Fold changes were calculated relative to control by the 2^−ΔΔCt^ method [30]. Statistical analysis was performed according to the comparative C_T_ method. All primer sets amplify a single product of the predicted size; primer sequences can be found in supplemental experimental procedures. β-actin was used as the constitutive control gene.

### Lentiviral shRNA infection and overexpression retrovirus infection

Cells were infected with either PLKO.1 non-targeting control lentivirus (Sigma) or shSTAT3 PLKO.1 lentivirus (clones TRCN0000020843 and TRCN0000020842), shJmjd3 (V2LHS_139678, Open Biosystems, Figure 2), shJmjd3 2 (pSico-R-PGKPuro lentivirus from [20], Supplementary Figure 2), in the presence of 8 μg/mL polybrene. 24 hours post-infection the virus containing media was removed and replaced with fresh stem cell media for 24 hours. Cells were then treated with 2.5 μg/mL puromycin. After 2-3 days of puromycin selection, cells were plated for assays as described in the figure legends.

The MSCV-Jmjd3 (Addgene plasmid 21212, [31]) MSCV-Jmjd3 mutant (Addgene plasmid 21214, [31]) and MSCV-Tap Control (Addgene plasmid 12570, [32] retroviral plasmids were obtained from Addgene (Cambridge, MA). The viruses were packaged in 293GPG cells as described by Ory et al [33]. Glioma stem cells were infected at an approximate MOI of 1 overnight, and then overnight again. 96 hours after the first infection cells were treated with 2.5 ug/mL puromycin for 4 days. After 4 days of puromycin selection cells were plated for assays as described in the figure legends.

### ChIP and ChIP-Sequencing

Cells were lysed, fixed, and sonicated according to the methods of De Santa et. al [34]. Sonicated lysates were incubated overnight with either anti-STAT3 (Santa Cruz Biotechnology, Santa Cruz, CA), anti-H3K27me3 (Millipore, Billerica, MA), or IgG control. Immunoprecipitation was carried out using StaphA cells according to the protocol of Kirmizis, Bartley (13) or using Protein A-Sepharose beads according to the nano ChIP-seq protocol (Adli and Bernstein (35)). After washing and cross-link reversal, immunoprecipitated DNA was purified using the Qiagen PCR Purification kit (Qiagen, Valencia, CA) and subjected to either qPCR or PCR as indicated in figure legends. For nano ChIP sequencing and nano ChIP-qPCR, the immunoprecipitated DNA was subjected to two intermediate rounds of amplification (Adli and Bernstein (35)) with custom primers containing barcodes that allowed us to multiplex 3 replicates for each of the two treatment groups in a sequencing lane (supplemental procedure). It was then subjected to either library amplification using TruSeq ChIP Sample preparation kit (Illumina Inc, San Diego, CA) followed by next generation sequencing on HiSeq 2500 (Illumina Inc, San Diego, CA) or qPCR. Myt1 primers were first described in [36]. FGF21 and GDF15 primers were designed to span regions of H3K27me3 enrichment near the gene promoter from ChIP-Sequencing data analysis. Primers used appear in supplemental experimental procedures. ChIP qPCR data was analyzed according to the percent of input method described by Haring et. al [37].

### ChIP-Sequencing data analysis

ChIP Sequencing reads were aligned to the human genome (hg19) using Bowtie in GALAXY platform and visualized in the UCSC genome browser [38]. The average read count through the 5kb upstream promoter region of each gene was computed across the genome. The normalized read counts at the promoters of STAT3 inhibited and control treatment groups was compared and a list of genes that showed reduction in H3K27me3 mark upon STAT3 inhibition was obtained [39, 40].

### Microarray

Microarray gene expression data was generated using GeneChip Human gene U133A 2.0 Arrays (Affymetrix, Santa Clara, CA). The CEL files were processed by Transcriptome Analysis Console (Affymetrix) software to analyze differentially expressed genes based on fold change.

## RESULTS

### STAT3 represses Jmjd3 expression in glioblastoma stem cells

We first examined Jmjd3 expression in response to STAT3 inhibition in two glioblastoma stem cell lines, GS6-22 and GS7-2, which we have previously characterized [11]. Upon STAT3 inhibition with the SH2-domain-targeting small molecule inhibitors STA-21 and S3I-201[41, 42] Jmjd3 expression is upregulated at both the protein and mRNA levels by 2-15 fold (Figure 1A, Figure S1A,B). Jmjd3 mRNA is also upregulated upon STAT3 knockdown using two different shRNAs (Figure 1B, Figure S1D-E). Additionally, Jmjd3 mRNA is upregulated by a highly potent next-generation STAT3 inhibitor BP-1-102 [43] and S3I-201 treatment in additional GBM-SC lines tested (Figure S1B). Thus, multiple STAT3 specific inhibitors lead to upregulation of Jmjd3 expression in different GBM-SC lines.

**Fig 1.**
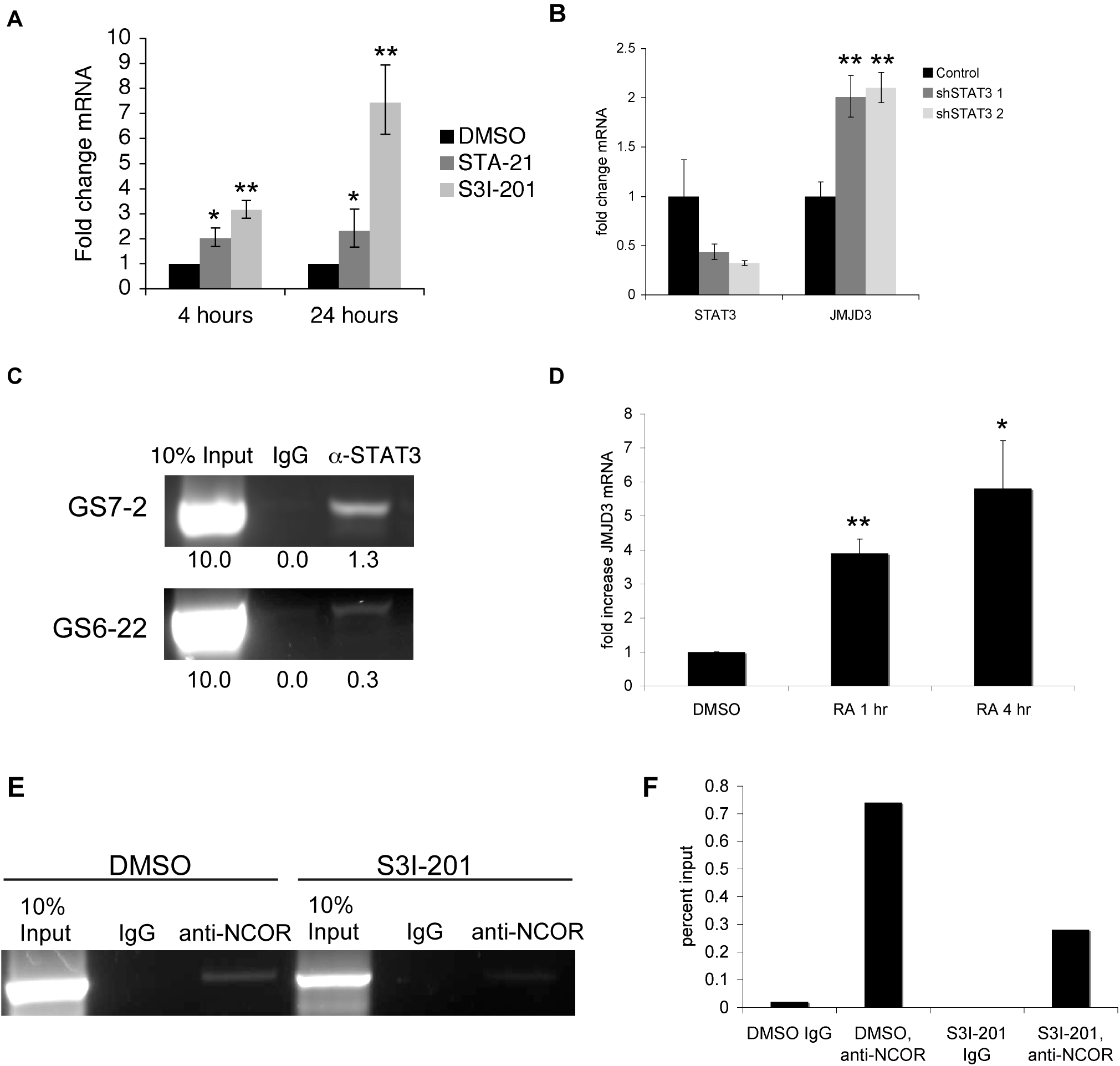
STAT3 represses Jmjd3 expression in glioblastoma stem cells. A. RT-qPCR of GS6-22 cells treated with S3I-201 or STA-21 demonstrates that Jmjd3 mRNA is upregulated at both 4 hours and 24 hours after inhibitor treatment. Values represent the fold change relative to DMSO treated cells for three experiments; bars SD (**p<0.01). B. Knockdown of STAT3 using an shRNA-containing lentivirus leads to the upregulation of Jmjd3 mRNA in GS6-22 cells, two days after selection. Values represent the fold change relative to DMSO treated cells for three experiments; bars SD (**p<0.01). C. Chromatin immunoprecipitation using anti-STAT3 antibody, followed by PCR using primers to the Jmjd3 promoter demonstrates STAT3 binding in GS7-2 and GS6-22 cells. Enrichment of DNA in STAT3 immunoprecipitation (relative to input) was quantified by ImageJ. D. Treatment of GS6-22 cells with retinoic acid increases Jmjd3 mRNA levels. GS6-22 cells were treated for 1 and 4 hours with 1 µM RA. Values represent the fold change relative to DMSO treated cells for three experiments; bars SD (*p<0.05, **p<0.01). E. S3I-201 treatment decreases NCor binding at the Jmjd3 promoter. GS6-22 cells were treated with S3I-201 (50 μM) or DMSO for 4 hours, fixed, lysed, and sonicated. Sonicated lysates were incubated overnight with anti-NCor or IgG control and subjected to chromatin immunoprecipitation. The resulting DNA was subjected to PCR using primers to the Jmjd3 promoter. F. Densitometry of E was performed using ImageJ.

By chromatin immunoprecipitation assay, STAT3 was found to bind to the Jmjd3 promoter at a position containing a conserved STAT3 binding site (Figure 1C, Figure S1C), suggesting that Jmjd3 is a direct STAT3 target gene. Previously it has been shown that Jmjd3 is upregulated during murine neural stem cell differentiation, and that it targets several neural specific genes upon differentiation of neural stem cells [19, 20]. Consistent with this we have found that Jmjd3 is upregulated during GBM-SC differentiation as well (Figure 1D, Figure S1F). UTX, another member of the Jmjd3 family of H3K27me3 demethylases, is not consistently upregulated by the two STAT3 inhibitors, S3I-201 and STA-21, in both of our cell lines (Figure S1G)

Interestingly, STAT3 appears to repress Jmjd3 expression, rather than activate it as is typical for STAT3 target genes [44]. However, STAT3 has been shown to repress genes in other cell types, and is associated with both transcriptionally active and repressed genes in embryonic stem cells [45] and in neural stem cells[46]. In murine neural stem cells, Jmjd3 is repressed by the NCoR and SMRT nuclear co-repressors, which associate with the retinoic acid receptor to repress transcription [19]. STAT3 has been shown to associate with the retinoic acid receptor (RAR), so we considered the possibility that NCoR repression is involved in regulating Jmjd3 in glioblastoma stem cells [47, 48]. Retinoic acid treatment of GS6-22 cells leads to a rapid increase in Jmjd3 mRNA (Figure 1D). Additionally, the NCoR co-repressor can be found at the Jmjd3 promoter by chromatin immunoprecipitation (Figure 1E,F). NCoR can be immunoprecipitated using the same primers as STAT3, suggesting it binds a similar region of the Jmjd3 promoter. Furthermore, treatment with the STAT3 inhibitor S3I-201 decreases NCoR binding to the Jmjd3 promoter (Figure 1E,F). It is possible, then, that STAT3 represses Jmjd3 expression by interaction with the NCoR-RAR complex, and that treatment with S3I-201 abolishes this interaction, thereby relieving the repression of Jmjd3.

### STAT3 regulates GBM-SC neurosphere formation and proliferation through repression of Jmjd3

Neurosphere formation is a hallmark of neural stem cells, and the ability to culture neurospheres from glioblastomas strongly correlates with poor patient prognosis [49]. Because STAT3 represses Jmjd3 expression in GBM-SC, we investigated whether knockdown of Jmjd3 expression could rescue the abrogation of neurosphere formation that accompanies STAT3 inhibition. GS6-22 and GS7-2 cells were infected with lentiviruses expressing shRNA to Jmjd3 or with control lentiviruses. Control infected cells exhibited decreased neurosphere formation in response to S3I-201 treatment as expected (Figure 2A,B,C). shJmjd3 infected cells, however, were able to form spheres in the presence of S3I-201 (Figure 2A,B,C). This was also true for cells infected with a second, distinct shRNA to Jmjd3, indicating that this is a specific effect of Jmjd3 knockdown (Figure S2A,B). This effect was more pronounced in the GS7-2 cells (Figure 2C) than the GS6-22 cells (Figure 2B) likely due to the enhanced degree of knockdown in the GS7-2 cells (78% knockdown in the GS7-2 cells versus 55% for GS6-22 cells) (Figure S2C,D). These data indicate that STAT3 repression of Jmjd3 is necessary for neurosphere formation.

**Fig 2.**
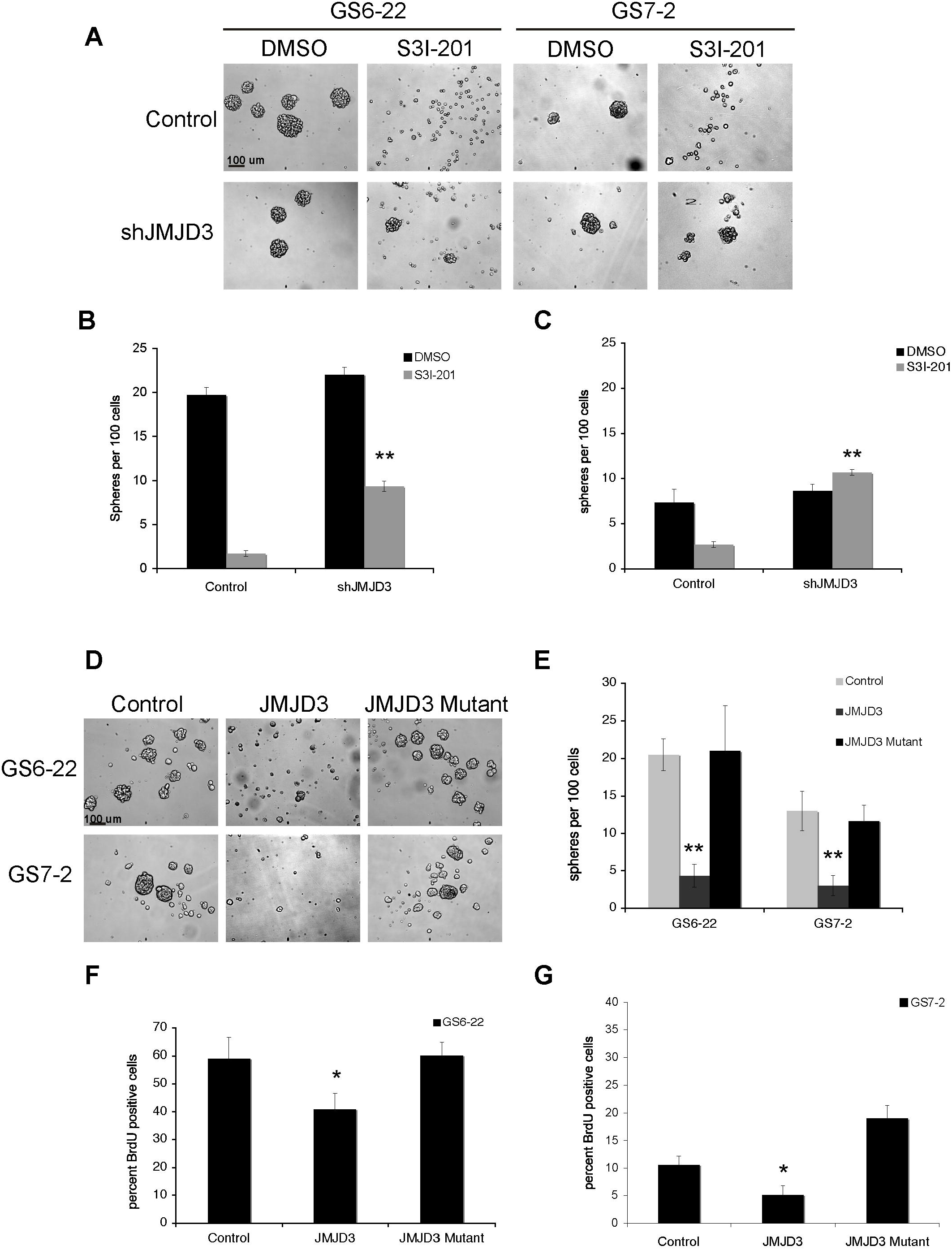
STAT3 controls GBM-SC neurosphere formation and proliferation through repression of Jmjd3. A. Knockdown of Jmjd3 rescues neurosphere formation in the presence of S3I-201. Representative images of GS6-22 and GS7-2 cells infected with either control or shJmjd3 containing lentivirus treated with DMSO or S3I-201 (50 μM). Images for GS6-22 cells were taken after 6 days and GS7-2 cells were taken after 4 days, based on differences in sphere formation rate in these lines. B. Quantification of neurosphere formation in GS6-22 cells. After 6 days of inhibitor treatment, the number of spheres per 100 cells was counted. Values represent the mean of triplicates within each treatment; bars SE (**p<0.01). C. After 4 days of inhibitor treatment, GS7-2 cell sphere formation was quantified as described for GS6-22 cells. Values represent the mean of triplicates within each treatment; bars SE (**p<0.01 relative to DMSO control, upon shJMJD3 infection). D. Representative images of GS6-22 and GS7-2 cells infected with overexpression retrovirus containing either a control (empty) plasmid, Jmjd3, or a catalytic domain mutant of Jmjd3. After three days of selection, cells were dissociated and replated at 100 cells per ml. Pictures were taken at 50X seven days after replating. E. Quantification of neurosphere formation capacity in GS6-22 and GS7-2 cells infected with the Jmjd3 overexpression retrovirus. Neurosphere formation assay was performed in triplicate (**p<0.01). F. GS6-22 and G. GS7-2 cells were infected with either control, Jmjd3, or Jmjd3 mutant retroviruses as previously described. Cells were pulsed with 30 μM BrdU for 16 hours. After 24 hours, cells were fixed and stained with an anti-BrdU antibody. Cells were also stained with 7-AAD at this time. The percentage of BrdU positive cells was analyzed using flow cytometry. Values represent the mean of 3 experiments; bars SD of the mean (*p<0.05).

Consistent with this, overexpression of Jmjd3 recapitulates significant aspects of the STAT3 inhibition phenotype. Both GS6-22 and GS7-2 cells that had been engineered to constitutively express Jmjd3 fail to form neurospheres in stem cell media (Figure 2D,E). Overexpression of a catalytic dead mutant of Jmjd3, however, does not inhibit sphere formation (Figure 2D,E). In addition, overexpression of Jmjd3, but not mutant Jmjd3, decreased proliferation in both GS6-22 and GS7-2 cells (Figure 2F,G). This effect was more pronounced in GS7-2 cells, but the degree of overexpression was also higher in these cells (Figure S2E,F). This indicates that the demethylase activity of Jmjd3 is necessary for the inhibition of sphere formation and proliferation of GBM-SC. STAT3 repression of Jmjd3, then, is required for GBM-SC neurosphere formation and proliferation.

Interestingly, overexpression of both wildtype Jmjd3 and the mutant increased apoptosis in GS6-22 cells (Figure S2G). It is possible the mutant Jmjd3 may interfere with some cellular functions by substrate binding and sequestration. Consistent with this, Jmjd3 has been found to have demethylase independent functions[50]. This effect was not observed in GS7-2 cells (Figure S2H). This may be because these cells lack expression of the INK4A/ARF locus (see below).

### STAT3 regulates H3K27 trimethylation and neural gene induction in GBM-SC

To identify putative STAT3 regulated genes that show induction of gene expression concurrent with histone H3K27 demethylation in GS6-22, we performed microarrays and genome wide CHIP-sequencing analysis using an antibody specific to H3K27me3, following STAT3 inhibition. Differential analysis of STAT3 inhibited and control GS6-22 cells yielded a ranked list of genes that were likely to show reduction in H3K27me3 mark (Table S1) or are induced upon STAT3 inhibition with a fold change >2 (Table S2). The intersected list of genes showing both reduction in repressive H3K27me3 mark as well as 2 fold or higher gene expression upon S3i treatment is tabulated in Table S3.

We subsequently confirmed that STAT3 inhibition by the next-generation STAT3 inhibitor BP-1-102 [43] upregulates mRNA expression of Fibroblast Growth Factor 21 (FGF21) and Growth Differentiation Factor 15 (GDF15) in GS6-22 and GS7-2 cells (Fig 3A, S3A). These genes are known to play roles in neural development [51, 52]. GDF15 has also been shown to induce apoptosis and inhibit tumorigenesis in GBM [53, 54].

**Fig 3.**
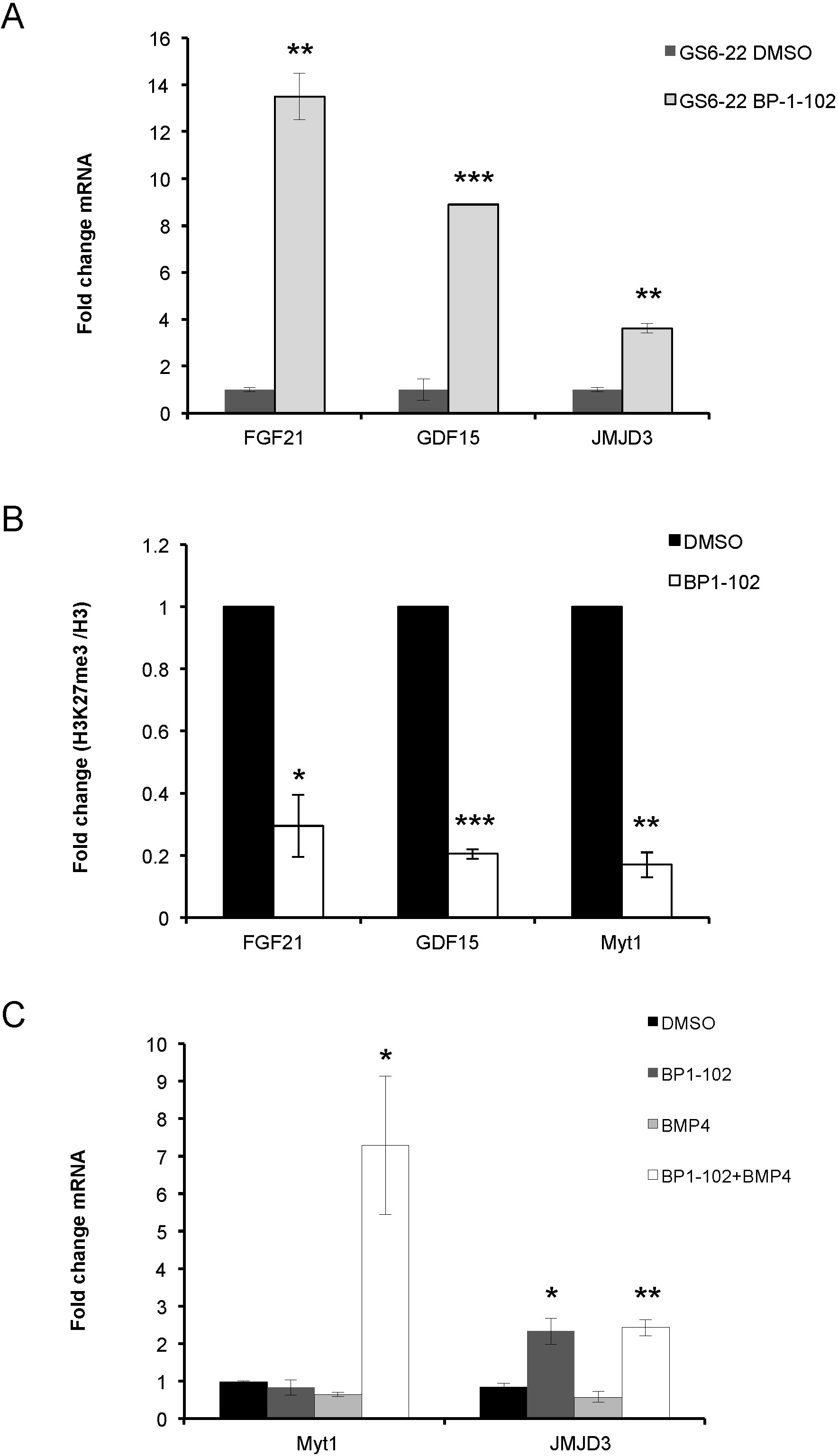
STAT3 regulates H3K27 trimethylation and neural gene induction in GBMSC. A. RT-qPCR analysis of FGF21, GDF15 and JMJD3 expression in GS6-22 cells treated with BP-1-102 (15uM) for 6 hours. Values represent the fold change relative to control cells for three experiments; bars SD (**p<0.01, ***p<0.005). B. GS6-22 cells were treated with BP-1-102 (15 μM) or DMSO for 6 hours, fixed, lysed, and subjected to chromatin immunoprecipitation with an antibody to H3K27 trimethylation or total histone H3. Quantitative PCR using primers to FGF21 or GDF15 or Myt1 genes was performed after immunoprecipitation, amplification and DNA purification. Data is displayed as percentage of input DNA of H3K27me3/H3 ChIP, normalized relative to control cells for three experiments (*p<0.05, **p<0.01, ***p<0.005). C. Demethylation of H3K27 at the Myt1 promoter confers BMP4 inducibility of the gene. RT-qPCR analysis of Myt1 and JMJD3 expression in GS6-22 cells treated with BP-1-102 (15uM), recombinant BMP4 (10 ng/ml) or a combination of both for 6 hours. Values represent the fold change relative to DMSO treated control cells for three experiments; bars SD (*p<0.05, **p<0.01).

Consistent with the induction of gene expression, ChIP-sequencing data analysis suggested loss of H3K27me3 peaks at FGF21 and GDF15 promoters in STAT3 inhibited GS6-22 cells compared to control. This was confirmed using ChIP-qPCR analysis, which showed that STAT3 inhibition by BP-1-102 decreases H3K27 trimethylation at FGF21 and GDF15 promoters (Fig 3B), concurrent with JMJD3 upregulation in GS6-22 cells (Fig 3A). The amount of total histone H3 does not change upon BP-1-102 treatment indicating that there is a loss of H3K27 trimethylation at these promoters and not just a loss of histone H3 association (Fig S3B, C). These results indicate that inhibition of STAT3 induces H3K27 demethylation and subsequent expression of target genes involved in neural differentiation of GBM-SC.

### Demethylation of H3K27 at the Myt1 promoter confers BMP4 inducibility

Because Jmjd3 inhibits sphere formation and proliferation of GBM-SC in a demethylase-dependent manner, we examined the H3K27me3 status at the promoter of a known Jmjd3 and polycomb target gene, myelin transcription factor 1, Myt1 [13], which is a gene that has been implicated in neuronal differentiation [55]. GS6-22 cells were treated with STAT3 inhibitor, BP-1-102, and subjected to chromatin immunoprecipitation (ChIP) using an antibody specific to H3K27me3. In GS6-22 cells, a decrease in H3K27 trimethylation was observed at the Myt1 promoter (Fig. 3B, S3D), concurrent with Jmjd3 upregulation. The degree of decrease in H3K27 trimethylation is consistent with that seen at other promoters demethylated by Jmjd3 [19, 56] and suggests that Myt1 is a Jmjd3 target gene in GBM-SC.

However, STAT3 inhibition or expression of Jmjd3 in stem cell growth conditions was not sufficient to induce Myt1 gene expression (Figure 3C, Fig. S4B), suggesting that other factors or signals are required to activate expression of these neural specific genes. It has been previously shown that Bone Morphogenetic Protein 4 (BMP4) activates the Smad signaling cascade and induces differentiation of stem-like, tumor-initiating precursors of GBMs [57] and that SMADs and Jmjd3 co-regulate genes in NSCs [58]. Myt1 expression is strongly induced upon BMP4 treatment when STAT3 is inhibited in GS6-22 and GS7-2 cells (Figure 3C, S4A). This suggests that STAT3 inhibition and subsequent demethylation of histone H3K27 at the promoter renders the chromatin accessible to BMP4 induced transcription factors at this locus.

Interestingly, we have not found a change in H3K27me3 at the *INK4A/Arf* locus (Figure S4C) in GS6-22 cells. This locus is a well-characterized Jmjd3 target [15, 56, 59], and its demethylation is necessary for the induction of growth arrest and senescence in several cell types. In GS7-2 cells, we failed to see PCR amplification of INK4A/ARF in genomic DNA, which suggests that this locus is deleted in GS7-2 cells (Figure S4D). This is not surprising given that over half of glioblastomas exhibit homozygous deletion of this locus [60]. Together, these observations suggest *INK4A/ARF* is not necessary for Jmjd3 regulation of GBM-SC proliferation and sphere formation. Ene et al (2012) came to a similar conclusion[29].

### STAT3 regulates Jmjd3 expression in human neural stem cells

Finally, we examined whether STAT3 repression of Jmjd3 was specific to glioblastoma stem cells, or whether the STAT3 inhibition phenotype is recapitulated in normal human neural stem cells. In neural stem cells derived from H9 embryonic stem cells [61, 62], STAT3 is activated by phosphorylation on both pTyr705 and pSer727 (Figure 4A). S3I-201 treatment of these cells inhibited neurosphere formation (Figure 4B) and lead to upregulation of Jmjd3 mRNA (Figure 4C), as well as to dose-dependent inhibition of proliferation as judged by BrdU incorporation (Figure 4D). STAT3, then, regulates neurosphere formation and proliferation in normal embryonic neural stem cells as well as in glioblastoma stem cells. STAT3 repression of Jmjd3 is also maintained in normal neural stem cells. This indicates that STAT3 control of the epigenetic program through Jmjd3 repression is a key mechanism of stem cell maintenance in both normal and tumorigenic neural stem cells.

**Fig 4.**
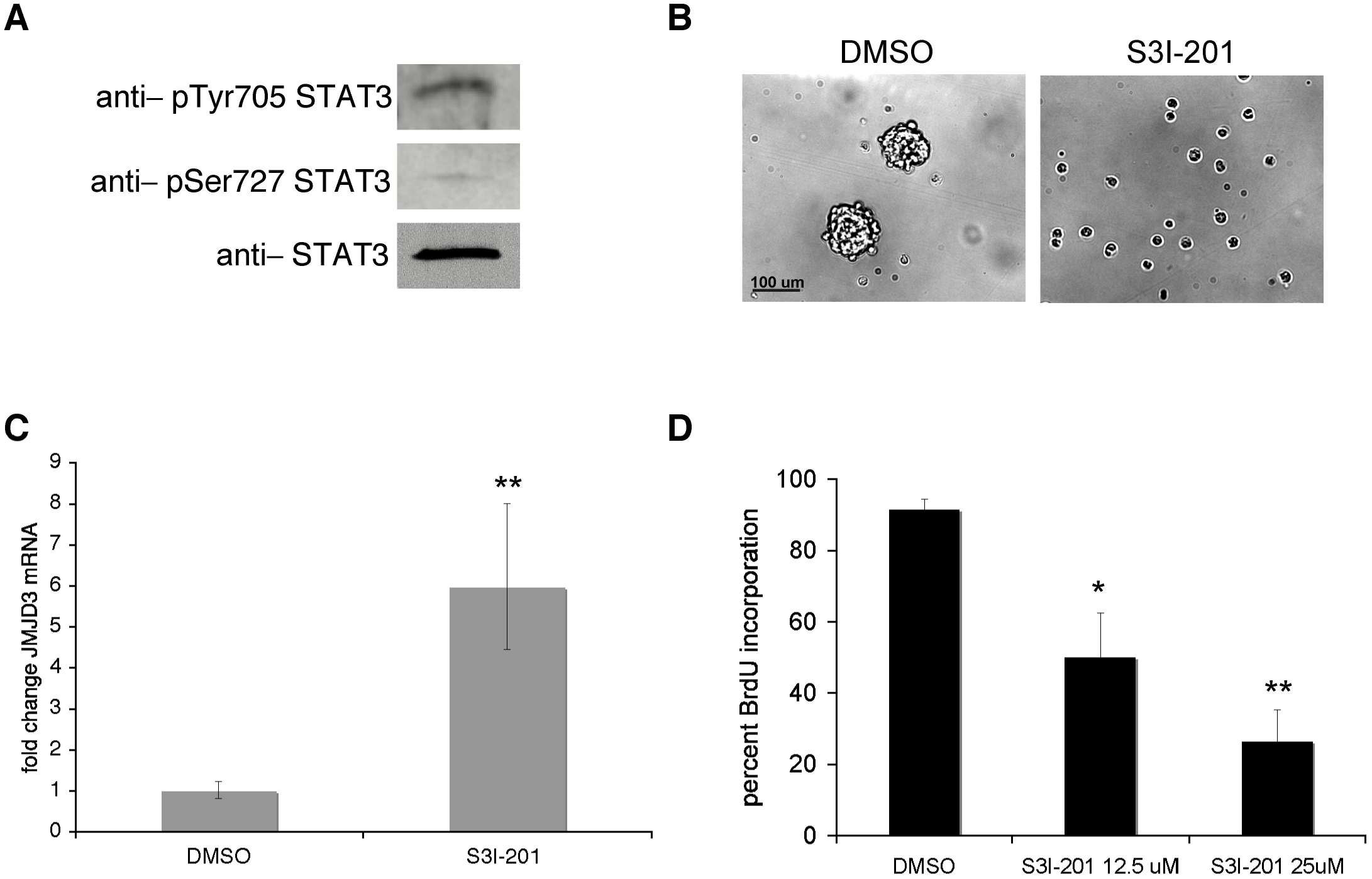
STAT3 inhibition in human neural stem cells leads to a decrease in neurosphere formation and proliferation, and an upregulation of Jmjd3 expression. A. STAT3 activation in human neural stem cells was assessed by immunoblotting with phospho-specific antibodies. B. Human neural stem cells were plated in serum free media supplemented with EGF and FGF2. 24 hours after plating, cells were treated with S3I-201 (50 μM) or DMSO control. Pictures were taken 3 days after treatment. C. STAT3 regulation of Jmjd3 was assessed by qRT-PCR. Human neural stem cells were treated with S3I-201 (50 μM) for 24 hours. Values represent the fold change relative to control cells for three experiments; bars SD (**p<0.01). D. Cells were treated with DMSO or S3I-201 for 24 hours, and then subjected to a 15 μM BrdU pulse for 20 hours. Values represent the average of three experiments; bars SE (*p<0.05, **p<0.01).

## Discussion

We have shown that STAT3 maintains normal neural and glioblastoma stem cells in a proliferative, self-renewing state via the repression of the histone demethylase Jmjd3. STAT3 repression of Jmjd3 is consistent with our previous data that STAT3 inhibition is irreversible in GBM-SC. We have found that STAT3 inhibition or knockdown leads to rapid upregulation of Jmjd3. We have also found that STAT3 binds to the Jmjd3 promoter in human GBM-SC which is consistent with published genomic data from murine ES cells [45] and suggests that STAT3 is a direct regulator of JMJD3.

While STAT3 is best known as a transcriptional activator, there is evidence that it can act as a repressor [63, 64]. ChIP-ChIP and microarray studies suggest that STAT3 represses genes in multiple cell types including in GBM-SC [45, 46, 65, 66] Previous studies indicated that NCoR negatively regulates Jmjd3 [19]. We have found that following STAT3 inhibition NcoR binding is reduced at the same promoter fragment that binds STAT3, suggesting that the mechanism of repression is recruitment or stabilization of the NCoR repressor complex at the promoter. While Ene et al (2012)[29] have found evidence that DNA methylation can also contribute to Jmjd3 repression, the sites of that STAT3 binds to appear to be distinct from the regions of DNA methylation [29]. However, given recent evidence that STAT3 can control DNA methylation [67], this mechanism of regulation merits further investigation.

We have also found that Jmjd3 overexpression inhibits neurosphere formation and cell proliferation (Figures 2,3). Consistent with our data, Jmjd3 overexpression has been shown to decrease the tumorigenicity of GBM-SC *in vivo* [29]. By contrast, another study reported that GSK-J4 mediated inhibition of JMJD3 and UTX was shown to reduce the sphere forming capacity of GBM-SC [68]. However, GSK-J4 has nonspecific inhibitory effects on other Jumonji domain family members[69], which need to be assessed to resolve such context-dependent effects. Moreover, targeted knockout of JMJD3 in the mouse brain indicates that expression of JMJD3 and its demethylase activity is needed for adult brain neurogenesis consistent with our results here[21].

We have found that overexpression of Jmjd3, which opposes PRC2 activity, also induces apoptosis in GS6-22 cells. A similar effect on both proliferation and apoptosis of GBM-SC has been reported for knockdown of BMI-1, part of the PRC1 complex [70]. Similarly, inhibition of EZH2, the enzymatic part of the PRC2 complex, causes a decrease in sphere formation and proliferation of GBM-SC [24]. Interestingly, EZH2 also methylates and regulates STAT3 in GBM-SC which further suggests a tight coordination between STAT3 and H3K27 trimethylation [71].

In this regard, it is also worth noting the role of histone H3K27 trimethylation in brain tumors is complex. While induction and maintenance of this mark is pro-oncogenic in medulloblastoma and adult glioblastomas[23–25, 71], global loss of H3K27 methylation is oncogenic in pediatric gliomas and in some adult GBMs[26, 27, 72]. Thus, the effects of H3K27 trimethylation on oncogenesis are context dependent. This is consistent with findings from other cancers as well [73, 74].

We have also found that a catalytic mutant of Jmjd3 does not affect neurosphere formation or proliferation. Thus, Jmjd3 upregulation and the subsequent demethylation of histone H3K27 on target genes is likely necessary for the decrease in GBM-SC growth. This induction of Jmjd3 likely imparts widespread changes to the chromatin and sustained mRNA expression of target genes involved in neural differentiation of GBM-SC. This would explain why either the restoration of STAT3 signaling following removal of S3I-201 STAT3 inhibitor [11], or sustained STAT3 signaling when Jmjd3 is constitutively expressed, does not rescue neurosphere formation. This difference in epigenetic state may also explain how STAT3 can induce astrocytic differentiation of neural stem cells under some conditions and preserve neural stem cell self-renewal in others [6, 75–77]. A similar persistence of a stable STAT3 phenotype has been demonstrated in breast epithelial cells, in which a transient expression of either of two STAT3-regulated microRNAs is sufficient to induce and maintain a stably transformed state that is also polycomb mediated [78, 79]. Thus, there is precedence for the transient upregulation of a single STAT3 target gene imparting permanent change to the cellular state.

Interestingly, although STAT3 inhibition induces neural genes, such as FGF21 and GDF15, neither Jmjd3 overexpression or STAT3 inhibition are sufficient to drive differentiation of the glioblastoma stem cells (Figure S4B, [11]). While Jmjd3 overexpression is sufficient to induce differentiation in some systems [19, 31], it is likely that additional positive differentiation signals are required both in these cases and in the case of the GBM-SC. Histone H3K27 demethylation may be sufficient to open the chromatin at specific genes, but positively acting transcription factors are likely needed in addition to bind to these promoters to activate transcription. Consistent with this, we have found that Jmjd3 overexpression can demethylate histone H3K27 at certain differentiation-specific genes such as Myt1, but the gene is upregulated only in the presence of recombinant BMP4. This situation is similar to previous results for the GFAP gene where epigenetic changes must occur before factors like CNTF are able to induce the gene during astrocytic differentiation [80–82]. Interestingly, STAT3 regulates GFAP as well [80–82].

In addition, Jmjd3 overexpression alone does not completely recapitulate the STAT3 inhibition phenotype, at least with respect to βIII-tubulin expression (Figure 3D, S4B, and [11]). Given that STAT3 signaling likely regulates many genes in addition to Jmjd3 in GBM-SC, this is not surprising. However, the fact that constitutive Jmjd3 expression substantially inhibits neurosphere formation as well as proliferation indicates that Jmjd3 repression by STAT3 is required for glioblastoma stem cell maintenance in the cell lines tested As STAT3 regulation of Jmjd3 was found in GBM-SC derived from multiple patient tumors, it is likely that this finding indicates a broad role for STAT3/Jmjd3 regulation of GBM-SC maintenance.

Strikingly, we have also found that STAT3 repression of Jmjd3 is a mechanism shared by both glioma stem cells and normal neural stem cells (Figure 4). While this could complicate therapeutic targeting of STAT3 in glioblastoma, it does provide further evidence that similar signaling and epigenetic mechanisms that govern both cancer stem cells and tissue stem cells at least in this case. STAT3 is necessary for the self-renewal of a variety of stem cell types, including murine embryonic stem cells [5], and its regulation of the epigenetic program may prove to be a widespread and developmentally critical mechanism of maintaining stem cell self-renewal.

## Acknowledgements

We thank G. Testa for shRNA to Jmjd3, Y. Shi for anti-Jmjd3 antibodies, P. Khavari for the Jmjd3 retroviral vectors obtained from Addgene, and H. Wakimoto for the MGG8 cell line. S3I-201 (NSC 74859) and STA-21 (NSC 628869) inhibitors were obtained from the NCI/DTP Open Chemical Repository (http://dtp.cancer.gov). The Tufts Neuroscience Core (P30 NS047243) provided the Stratagene qPCR machine. We thank J. Wu and A. Yee for critical reading of the manuscript. This work was supported in part by NIH R01NS072414 to BHC.

## Supporting Information

S1 Fig. STAT3 represses JMJD3 expression in glioblastoma stem cells.

S2 Fig. JMJD3 knockdown partially rescues neurosphere formation in S3I-201.

S3 Fig. Neural gene induction and H3K27 demethylation in BP-1-102treated GBM-SC.

S4 Fig. MYT1 expression and INK4A/ARF H3K27 methylation in GBM-SC.

S1 Table. GS6-22 cells were treated with S3I-201 (50uM) or DMSO control for 3 days. ChIP Sequencing analysis using H3K27me3 antibody yielded a list of gene promoters that likely show a reduction in repressive H3K27me3 mark upon S3i treatment.

S2 Table. GS6-22 cells were treated with S3I-201 (50uM) or DMSO control for 3 days. Microarray analysis yielded a list of genes that likely show 2 fold or higher gene expression upon S3i treatment.

S3 Table. GS6-22 cells were treated with S3I-201 (50uM) or DMSO control for 3 days. Intersected list of genes showing both reduction in repressive H3K27me3 mark as well as 2 fold or higher gene expression upon S3i treatment.

S1 File Supplemental Experimental Procedures

